# STACAS: Sub-Type Anchor Correction for Alignment in Seurat to integrate single-cell RNA-seq data

**DOI:** 10.1101/2020.06.15.152306

**Authors:** Massimo Andreatta, Santiago J. Carmona

## Abstract

Computational tools for the integration of single-cell transcriptomics data are designed to correct batch effects between technical replicates or different technologies applied to the same population of cells. However, they have inherent limitations when applied to heterogeneous sets of data with moderate overlap in cell states or sub-types. STACAS is a package for the identification of integration anchors in the Seurat environment, optimized for the integration of datasets that share only a subset of cell types. We demonstrate that by *i)* correcting batch effects while preserving relevant biological variability across datasets, *ii)* filtering aberrant integration anchors with a quantitative distance measure, and *iii)* constructing optimal guide trees for integration, STACAS can accurately align scRNA-seq datasets composed of only partially overlapping cell populations. We anticipate that the algorithm will be a useful tool for the construction of comprehensive single-cell atlases by integration of the growing amount of single-cell data becoming available in public repositories.

**Code availability:** **R package:** https://github.com/carmonalab/STACAS

**Docker image:** https://hub.docker.com/repository/docker/mandrea1/stacas_demo

## 1 Introduction

Massively parallel single-cell transcriptomics (scRNA-seq) has emerged as a transformative technology that enables measuring molecular profiles at single-cell resolution. However, despite the highly multiplexed technologies, single-cell data are produced separately for different tissues and organs and are affected by multiple batch effects, such as different sample processing and scRNA-seq protocols. As such, integration of single-cell data might be the ultimate challenge in the field towards the generation of single-cell atlases (1–3).

Seurat (4) is currently one of the most popular and best performing algorithms for single-cell data integration, and can be effortlessly integrated into complex analysis pipelines (5). At the core of the Seurat integration algorithm is the identification of mutual nearest neighbors (MNN) across single cell datasets, named “anchors”, in a reduced space obtained from canonical correlation analysis (CCA). These anchors and their scores are used to compute correction vectors for each query cell, transforming (i.e. batch-correcting) its expression profile (6). Transformed cell profiles can then be jointly analyzed as part of an integrated space. To handle more than two datasets, a guide tree based on pairwise batch similarities is used to dictate the batch integration order. While Seurat has proven very powerful for the removal of technical artifacts between replicated experiments or even different sequencing technologies (5), it tends to overcorrect batch effects and performs poorly when integrating heterogeneous datasets (7), where only a fraction of cell types are shared between individual samples. This is crucial for the creation of reference cell type-specific single-cell atlases where the datasets to integrate were obtained from different tissues or experimental conditions (e.g. T cells from blood vs tumor-infiltrating T cells), and as a consequence are composed of different, partially overlapping cell states or sub-types.

## 2 RESULTS

STACAS is a package for determining integration anchors between heterogeneous datasets, and it is designed to be easily incorporated into Seurat dataset integration pipelines. STACAS employs a reciprocal principal component analysis (PCA) procedure to calculate anchors, where each dataset in a pair is projected onto the reduced PCA space of the other dataset; mutual nearest neighbors are then calculated in these reduced spaces. Crucially, and in contrast to the CCA reduction used by Seurat, the expression values of genes used in generating the PCA spaces are not rescaled to have zero mean and unit variance. When integrating heterogeneous datasets, for instance composed only of CD4+ or CD8+ cells, such rescaling can cancel out important biological differences between the datasets (**Figure 1A**).

**Figure 1:**
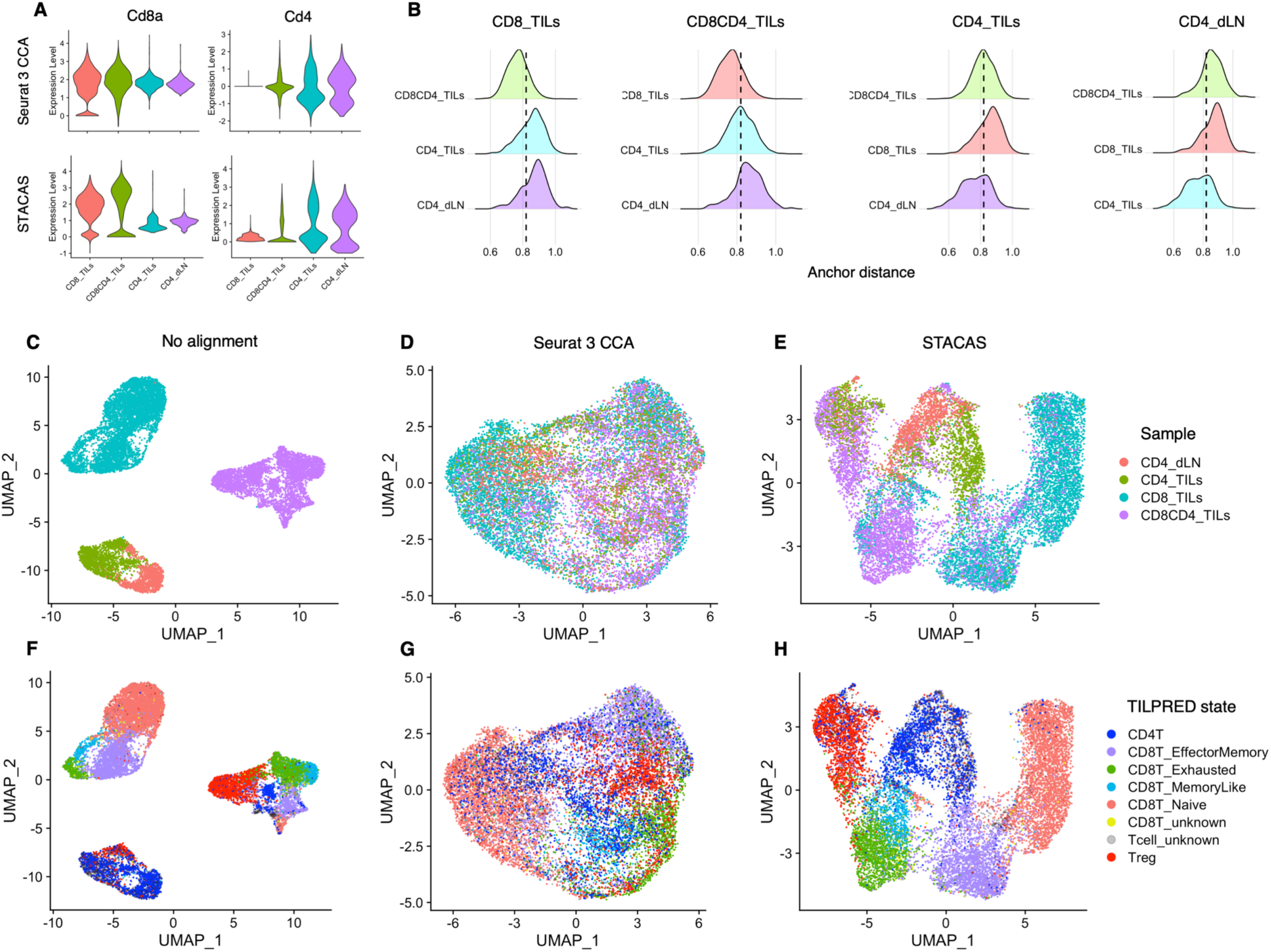
Anchor finding and dataset integration using STACAS. **A**) Expression level (*log* [normalized UMI counts + 1]) of *Cd8a* and *Cd4* after integration with Seurat CCA (top) or STACAS (bottom); important biological differences between the samples are lost by data rescaling and sub-optimal anchoring by Seurat 3 CCA. **B**) Anchor distance distribution between pairs of samples prior to anchor filtering by STACAS; poor anchors with distance higher than threshold (represented with a vertical dashed line) are filtered out by STACAS. **C-E)** Low-dimensionality UMAP visualization of scRNA-seq data, colored by sample, without batch correction (**C**), using Seurat CCA anchors (**D**) and using STACAS anchors (**E**) for dataset alignment. **F-H)** UMAP visualization of scRNA-seq data, colored by TILPRED state prediction, without batch correction (**F**), using Seurat CCA anchors (**G**) and using STACAS anchors (**H**) for dataset alignment.

A second innovation introduced in STACAS is the filtering of anchors based on anchor pairwise distance, which is calculated on the reduced PCA spaces used to determine the anchors. We observed that the distribution of anchor distances between datasets with shared cell subtypes (i.e. a CD4+CD8+ sample with a CD8+ sample) is centered on lower pairwise anchor distances compared with dataset pairs with limited or no overlap (e.g. a CD4+ sample and a CD8+ sample) (**Figure 1B**); anchor distance can then be used as a quantitative measure to filter spurious anchors and improve dataset integration. In STACAS, the anchor filtering threshold defaults to the 80th percentile of the distance distribution between the two most similar datasets.

Finally, the anchors determined by STACAS can be used directly for dataset integration using the *IntegrateData* function in Seurat 3. STACAS suggests a guide tree to determine the order in which datasets are to be integrated. In contrast to the Seurat default guide tree, which favors datasets with the highest total number of cells in any given pair, STACAS prioritizes samples with the highest total number of anchors; the rationale being that datasets with many anchors are likely to contain more cell types and represent the “centroid” of the integrated map.

In the example in **Figure 1**, we integrated four scRNA-seq datasets of mouse T cells from public repositories, composed of *i)* CD8+ tumor-infiltrating lymphocytes (TILs) (8); *ii)* CD4+ and CD8+ TILs (9); *iii)* CD4+ T cells from tumors (10); and *iv)* CD4+ T cells from draining lymph nodes (dLN) (10). There is an evident batch effect between the samples, with the cells of each sample clustering together regardless of their type (**Figure 1C** and **1F**). Consistently with a recent benchmark (7), dataset alignment using Seurat 3 appears to overcorrect these batch effects, overlaying samples with little in common such as CD4+ dLN and CD8+ TILs (**Figure 1D** and **1G**). In contrast, STACAS only aligns cells with similar states across samples, limiting the superposition of CD4+ with CD8+ cells (**Figure 1E**). Supervised cell state classification using TILPRED (8) confirms that in most cases STACAS was able to cluster cell types across different, heterogeneous data sets (**Figure 1H**). We obtained similar results on larger scale integration tasks towards the construction of reference T cell maps in cancer and chronic infection [manuscript in preparation].

STACAS is available as an R package at https://github.com/carmonalab/STACAS and as a Docker image, and can be easily incorporated in Seurat 3 pipelines for data integration. A demo detailing the functions and usage of the package with sample data can be found at https://gitlab.unil.ch/carmona/STACAS.demo

## Notes

https://github.com/carmonalab/STACAS

### Competing Interest Statement

The authors have declared no competing interest.

